# The effects of host species and sexual dimorphism differ among root, leaf and flower microbiomes of wild strawberries *in situ*

**DOI:** 10.1101/225755

**Authors:** Na Wei, Tia-Lynn Ashman

## Abstract

Plant-associated microbiomes profoundly influence host interactions with below- and aboveground environments. Characterizing plant-associated microbiomes in experimental settings have revealed important drivers of microbiota assemblies within host species. However, it remains unclear how important these individual drivers (e.g., organ type, host species, host sexual phenotype) are in structuring the patterns of plant–microbiota association in the wild.

Using 16s rRNA sequencing, we characterized root, leaf and flower microbiomes in three closely related, sexually polymorphic *Fragaria* species, in the broadly sympatric portion of their native ranges in Oregon, USA. Taking into account the potential influence of broad-scale abiotic environments, we quantified the relative effects of organ type, host species and sex on the α- and β-diversity of bacterial communities in these wild populations. Our statistical models showed that organ type explained the largest variation of compositional and phylogenetic α- and β-diversity of plant microbiomes, and its overall effect exceeded that of host plant species. Yet, the influence of host species increased from root to leaf to flower microbiomes. We found strong sexual dimorphism in flower and leaf microbiomes, especially in host species with the most complete separation of sexes. Our results provide the first demonstration of enhanced effects of host species and sexual dimorphism from root to flower microbiomes. While these findings await manipulative experiments to disentangle contributions of micro-environment and the residing microbiota from host genetic effects, we anticipate that such phenotypic patterns of host species–microbiota association may be broadly applicable to other plants in the wild.

## INTRODUCTION

The plant-associated microbiome is considered part of the extended plant phenotype (Bordenstein and Theis, 2015; Müller *et al*., 2016). Microbiota living on the surface and inside of host plants mediate processes vital to plant fitness, ranging from nutrient acquisition and stress responses, to pollination (Vorholt, 2012; Bulgarelli *et al*., 2013; Aleklett *et al*., 2014; Müller *et al*., 2016). Recent quantitative characterizations of plant-associated microbiomes have focused on microbiota harbored by leaves and roots in greenhouse and common garden settings (e.g., Lundberg *et al*., 2012; Peiffer *et al*., 2013; Bai *et al*., 2015; Edwards *et al*., 2015; Wagner *et al*., 2016), and have identified important drivers underlying microbiota assemblies in plants aboveground (AG) and belowground (BG).

These drivers include environment-related factors such as soil source (Edwards *et al*., 2015) and experimental site (Peiffer *et al*., 2013), and host-related factors including organ type (Bai *et al*., 2015), within-organ compartment (Lundberg *et al*., 2012), host species (Schlaeppi *et al*., 2014) and genotype (Wagner *et al*., 2016). However, comparative data are lacking that broaden AG and BG microbiomes to also encompass those associated with plant reproductive organs (e.g., flowers) and sexual phenotype, and generalize relevant findings from a single to multiple host species, especially in the wild. These are, nevertheless, an important first step towards understanding how plants recruit microbiota that may in turn affect host fitness in wild populations, and for determining whether individual drivers identified under experimental settings are also of paramount importance in the wild.

While consensus from studies in experimental or managed field settings is growing that organ type (root, leaf, flower; e.g., Zarraonaindia *et al*., 2015) and host plant species (Schlaeppi *et al*., 2014) influence microbial communities, their relative roles have not been explicitly quantified. This is in part because empirical studies profiling plant AG (leaf and flower) and BG (root) microbiomes are often not directly comparable, and vary in terms of environmental settings, microbial community type and anatomical details of plant organs. For example, comparative investigations of leaf microbiomes tend to focus on epiphytic microbiota, and have often involved tens to hundreds of host species in semi-natural or natural habitats (Redford *et al*., 2010; Kembel *et al*., 2014). By contrast, those on root microbiomes have typically evaluated microbiota of a few host species (Schlaeppi *et al*., 2014), or different genotypes and root compartments within a single host species (Lundberg *et al*., 2012; Peiffer *et al*., 2013; Edwards *et al*., 2015), often in experimental and/or managed field sites. Studies of flower microbiomes mostly focus on compartment differences, and/or in comparison to other organs, within a single host species in the wild or managed field sites (Ottesen *et al*., 2013; Junker and Keller, 2015; Zarraonaindia *et al*., 2015). While these studies begin to suggest a defining role of organ type in structuring bacterial microbiota in plants (Müller *et al*., 2016), the differences among studies make the quantitative inference of respective effects difficult. Quantifying microbiota across multiple organ types (root, leaf, flower) and host plant species *in situ* will provide a direct evaluation of the extent to which organ type exceeds host species in shaping microbial communities in the wild.

The effect of host species on microbiomes has been examined in controlled settings (in roots; Schlaeppi *et al*., 2014) and in the wild (in roots and/or leaves; e.g., Kembel *et al*., 2014; David *et al*., 2016). In wild populations, the host species–microbiota association can be influenced by range differences among plant species (e.g., Coleman-Derr *et al*., 2016). This is because the regional species pool of microbiota that potentially colonize host plants (e.g., those harbored by soil) can vary with broad-scale abiotic environments of plant species (de Vries *et al*., 2012). Additionally, even when ranges overlap and plants occur in sympatry, species can still vary in their microhabitats, where they also alter the local species pool of microbiota (Burns *et al*., 2015), by enriching and depleting certain microbes. As a result, host species effect on microbiomes in the wild could be viewed as (residual) host effect, after accounting for that of broad-scale abiotic environments (e.g., temperature, precipitation; Zimmerman and Vitousek, 2012). Such (residual) host species effect represents species–microbiota association attributable to host phenotype (i.e., the outcome of genotype and micro-environment), and micro-environment including local microbiota that are influenced by plant species; this is, by definition, the extended phenotype of host plants. Thus, we may expect host species effect on microbial communities to be stronger in the wild than in controlled settings, the knowledge of which is essential for understanding the variation of plant–microbiota association that can influence plant fitness in nature.

Microbiomes can be also influenced by host sex, as demonstrated in animal systems including humans (Markle *et al*., 2013; Dominianni *et al*., 2015). Such sexual dimorphism, however, has rarely been explored in plants (but see Golonka and Vilgalys, 2013), despite the fact that sexual phenotype (female, male or hermaphrodite) is known to influence floral and functional traits, and several ecophysiological processes (Barrett and Hough, 2013; Vega-Frutis *et al*., 2013). As a result, plant sexes may differ in the principles governing microbiota assemblies (i.e., dispersal, habitat filtering and niche partitioning). First, the species pool of colonizing microbes can vary between female and male plants, owing to differential visitation of pollinators that may carry and disperse microbes, as an outcome of differences in floral rewards and attractive traits between sexes (Ashman *et al*., 2000; Ashman *et al*., 2005). Second, plant sex can influence the niche space available for microbiota. Sexual dimorphism in flower size and longevity (Barrett and Hough, 2013) likely affects the size and dynamics of microbial habitats. Likewise, sexual dimorphism in leaf traits (e.g., trichomes and leaf toughness; Cornelissen and Stiling, 2005) can potentially define the living environments of leaf microbiota (Krimm *et al*., 2005; Vorholt, 2012). Finally, sex-differential susceptibility and/or allocation to defense (Kaltz and Shykoff, 2001; Ashman, 2002; Vega-Frutis *et al*., 2013) could alter resident microbial communities via microbe–microbe interactions (Müller *et al*., 2016). Comparisons across AG and BG organs would inform broadly on the potential for sexual differences in microbiomes.

In this study, we aim to quantify root, leaf and flower microbiomes in three closely related wild strawberries in the broadly sympatric portion of their native ranges. These perennial, sexually polymorphic *Fragaria* (Liston *et al*., 2014) include two octoploid species *F. chiloensis* and *F. virginiana* ssp. *platypetala*, and their natural hybrid (*F. ×ananassa* ssp. *cuneifolia*), which is the wild relative of the cultivated strawberry (*F. ×ananassa* ssp. *ananassa*). Specifically, we aim to address three key questions concerning the effects of host species, organ type and sexual phenotype in these wild populations: 1) What is the relative importance of host species and organ type in structuring microbial communities? 2) Does the strength of host species influence vary among root, leaf and flower microbiomes? 3) Do microbiomes differ between host plants of different sexual phenotype?

## MATERIALS AND METHODS

### *Fragaria* microbiome collection

Dioecious (male and female) *F. chiloensis* is a coastal specialist growing in front dunes, the soils of which are high in sand and low in nutrients, compared to fertile mesic forest-edge populations of subdioecious (hermaphrodite, male and female) *F. virginiana* ssp. *platypetala*, and the intermediate habitats of subdioecious *F. ×ananassa* ssp. *cuneifolia* (Salamone *et al*., 2013). These three species are widely distributed in western North America (Staudt, 1999). In Oregon, where they occur in sympatry but not in the same microhabitats (Salamone *et al*., 2013), we collected microbiota samples from seven wild populations (Figure 1A) over a 6-day period in May 2016: two populations of *F. chiloensis* (Salishan, ‘SAL’: 44.919° N, 124.027° W; Strawberry Hill, ‘SH’: 44.254° N, 124.112° W); two of *F. virginiana* ssp. *platypetala* (Willamette National Forest, ‘WNF’: 44.638° N, 121.941° W; Fisherman's Bend Recreation, ‘FBR’: 44.755° N, 122.515° W); three of the natural hybrid *F. ×ananassa* ssp. *cuneifolia* (Marys Peak, ‘MP1’: 44.497° N, 123.546° W; ‘MP2’: 44.507° N, 123.569–123.579° W; Corvallis, ‘COR’: 44.506° N, 123.285° W). From each population, we randomly selected two female and two male-fertile (male or hermaphrodite) plants that were at least 2 m apart from each other, and collected root, leaf and flower samples from each plant. However, at COR we only sampled roots and leaves from three plants, as this population passed flowering. In total, our collection comprised 78 samples for the three species with organ type of root (*N* = 27), leaf (*N* = 27) and flower (*N* = 24).

**FIGURE 1.**
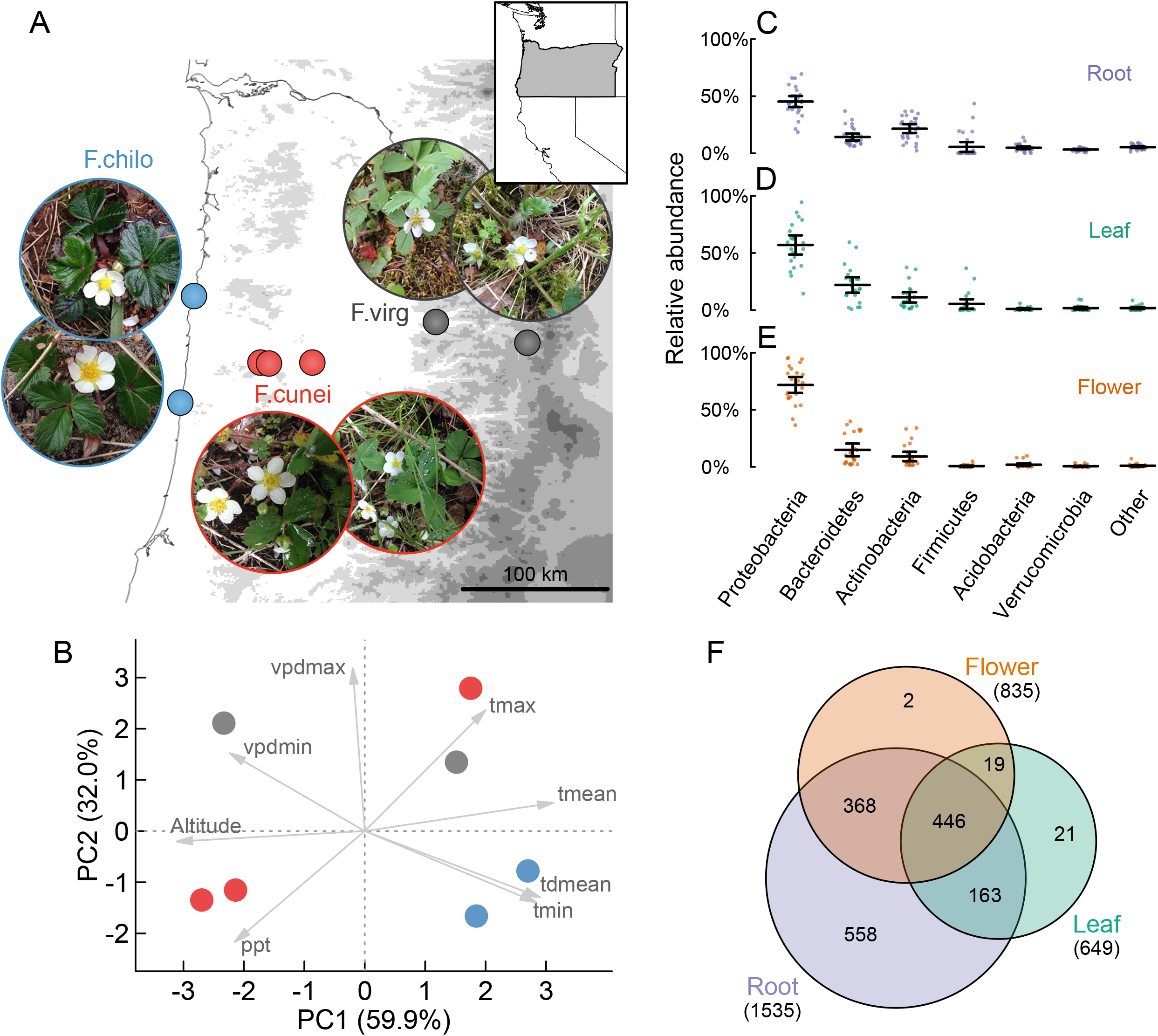
Collection and summary of *Fragaria* microbiomes. (A) Field collection of root, leaf and flower microbiomes of *F. chiloensis* (F.chilo), *F. virginiana* ssp. *platypetala* (F.virg) and *F. ×ananassa* ssp. *cuneifolia* (F.cunei) from seven wild populations (solid circles) in Oregon, USA. (B) PCA of climatic variables and elevation data of the sampled populations. The seven climatic variables include temperature (mean, tmean; minimum, tmin; maximum, tmax), mean dewpoint temperature (tdmean), precipitation (ppt) and vapor pressure deficit (minimum, vpdmin; maximum, vpdmin). The first two principal components visualized here (PC1 and PC2, denoted as PC1.clim and PC2.clim hereafter and in the main text) were used in statistical models to control for the effects of abiotic environments. (C–E) The relative abundances of major bacterial phyla in root, leaf and flower microbiomes, respectively. Dots represent individual microbial communities; means and error bars (2 s.e.) are indicated. (F) OTU overlap among root, leaf and flower microbiomes of all three host species.

For each plant, one flower (~2.5 cm in diameter), and the central leaflet (~2 cm in width) of a healthy trifoliate leaf that showed no evidence of herbivory and pathogen infection, were collected separately using ethanol-rinsed forceps and put directly into 750 μL Xpedition Lysis/Stabilization Solution in a ZR BashingBead Lysis tube (Zymo Research, Irvine, CA). Roots of the same plant were unearthed by the assisting person with ethanol-rinsed gloves, and were shaken vigorously to remove attached soil. Then five segments (~5 cm in length each) of fine roots, including some rhizosphere soil particles, were severed using sterile forceps and stored in the same manner. These samples were transferred to a −20 °C freezer within six hours after field collection, and shipped with dry ice to the University of Pittsburgh for DNA extraction.

Our leaf and flower samples contained both epiphytic and endophytic microbiota. The root samples also included some rhizosphere microbiota, in addition to rhizoplane and endosphere microbiota. For simplicity, we refer to these organ-associated microbiomes as root, leaf and flower microbiomes.

### DNA extraction and 16S rRNA amplicon sequencing

Samples were homogenized using a TissueLyser II (QIAGEN, Germantown, MD), and genomic DNA was extracted using Xpedition Fungal/Bacterial DNA MiniPrep (Zymo Research) under a sterile laminar flow hood. The same extraction procedure was conducted on three negative controls without plant samples. The amplification and sequencing of the V4 region of the 16S small subunit ribosomal RNA (rRNA) gene were performed at the Environmental Sample Preparation and Sequencing Facility at Argonne National Laboratory. In brief, the V4 region was amplified using 515F-806R primer pair (515F: GTGCCAGCMGCCGCGGTAA; 806R: GGACTACHVGGGTWTCTAAT) with 12-bp barcodes. Peptide nucleic acid (PNA) clamps designed from *Arabidopsis thaliana* (Lundberg *et al*., 2013) were added in amplification to reduce *Fragaria* plastid contamination. The three negative controls failed in PCRs and generated primarily primer dimers. The 16S rRNA amplicons of the 78 samples were sequenced using a 1/5 lane of 2 × 151 bp on an Illumina MiSeq instrument.

### OTU profiling and filtering

Paired-end reads were first joined using PEAR v0.9.6 (Zhang *et al*., 2014) with an overlap size of ≥20 bp. The successfully merged reads were used for subsequent open-reference operational taxonomic unit (OTU) picking. Sequence demultiplexing and quality filtering (with Phred quality scores of ≥20) were performed using QIIME v1.9.1 (Caporaso *et al*., 2010). The resulting sequences were clustered into OTUs based on a similarity threshold of ≥97% by PyNAST and assigned with taxonomic identification by RDP classifier based on the Greengenes reference database (13_8 release), as implemented in QIIME. After chimera removal using QIIME ChimeraSlayer, aligned OTU representative sequences were used to build a midpoint-rooted phylogenetic tree of these OTUs using QIIME FastTree.

The QIIME-generated OTU table was further filtered before the conversion into a microbial community matrix. First, we filtered out OTUs belonging to chloroplasts and mitochondria. Second, we removed the singletons as well as low-frequency OTUs that accounted for ≤0.01% of the total observations of the entire OTU table. Third, we removed low-depth samples of <100 observations (*N* = 6, five leaf and one flower samples). Fourth, we normalized the OTU table to the same observations for each sample using the median per-sample depth (of 2796), while keeping per-sample OTU relative abundances unchanged (McMurdie and Holmes, 2014). The resultant normalized OTU table was used as the microbial community matrix for downstream statistical analyses, because normalization using alternative per-sample depths (e.g., mean or maximum depth) and raw OTU table did not affect the results (data not shown).

### Abiotic environments of sampled populations

To account for abiotic effects on microbial communities, we used seven PRISM climatic variables (PRISM Climate Group, 2004) of the current (1981–2010) conditions at 30-arcsec resolution, and elevation data, for the seven sampled populations. The seven annual climatic variables include temperature (mean, minimum and maximum), mean dewpoint temperature, precipitation and vapor pressure deficit (minimum and maximum). We conducted a principal component analysis (PCA) of these variables, including elevation, using prcomp() in R v3.3.3 (R Core Team, 2017). The first two principal components (denoted as PC1.clim and PC2.clim) were taken as the abiotic predictors in the following statistical models.

### Statistical analyses of microbial community α-diversity

Species and phylogenetic α-diversity metrics considered Shannon diversity, Faith’s phylogenetic diversity (PD) and abundance-weighted mean phylogenetic distance (MPD), which were calculated using the R package vegan (Oksanen *et al*., 2017) and picante (Kembel *et al*., 2010). These α-diversity metrics were transformed (i.e., log_e_(PD), MPD^3^) to improve normality, and used as response variables in general linear mixed models (LMMs) using the package lme4 (Bates *et al*., 2015). The fixed effects included PC1.clim + PC2.clim + Species + Sex + Organ + Species:Sex + Species:Organ + Sex:Organ; the random effect included individual plants. We did not include populations in random effects for two reasons: first, models that incorporated nested random effects failed to converge given the sample size; second, individuals also captured some of the population variation. For the main effect of each predictor and their interactions, the least-squares means (LS-means) were estimated using the package lmerTest (Kuznetsova *et al*., 2016), and the statistical significance was evaluated by Type III sums of squares (SS). When considering organ type separately, we subdivided the microbial community matrix by organ type and re-estimated α-diversity metrics for organ-specific microbial communities. General linear models (LMs) were fitted with PC1.clim + PC2.clim + Species + Sex + Species:Sex, in which the LS-means and Type III SS were estimated using the package phia (Rosario-Martinez, 2015) and car (Fox and Weisberg, 2011), respectively.

### Statistical analyses of microbial community β-diversity

Compositional and phylogenetic β-diversity metrics considered Bray–Curtis dissimilarity (in vegan), inter-community MPD (betaMPD in picante) and abundance-weighted UniFrac distance in the package GUniFrac (Chen, 2012). Visualization of β-diversity metrics used the nonmetric multidimensional scaling (NMDS) in vegan for Bray–Curtis dissimilarity, and principal coordinates analyses (PCoAs) by cmdscale() for UniFrac distance and betaMPD.

These β-diversity metrics were taken as response variables in permutational multivariate analyses of variance (PERMANOVA) using vegan adonis2(). To assess the statistical significance (i.e., the marginal, instead of sequential, effect) of each main effect (or main term), PERMANOVA included PC1.clim + PC2.clim + Species + Sex + Organ. To assess the marginal effect for each interaction term, PERMANOVA included both the above main effects and their interaction terms (Species:Sex + Species:Organ + Sex:Organ). As a complement to PERMANOVA, generalized linear models (GLMs) with negative binomial errors were conducted using the package mvabund (Wang *et al*., 2012) to assess how multivariate community structures changed in response to the main and interaction terms. The marginal effect of each term was assessed by nested model comparison between a full model and a reduced model with the focal term removed using a likelihood ratio test. OTUs that responded significantly to each model term were identified using univariate likelihood ratio tests with *P* values adjusted by resampling-based multiple testing implemented in mvabund to control for the family-wise error rate (FWER; alpha = 0.05). Here we referred to mvabund GLMs as FWER-GLMs.

PERMANOVA and FWER-GLMs were also conducted to model microbial community β-diversity for each organ type separately. Visualization of organ-specific β-diversity metrics was performed using constrained PCoAs by vegan capscale().

### Differentially abundant OTUs among microbial communities

As a complement to the univariate tests of individual OTU abundances in mvabund, we used the package edgeR (Robinson *et al*., 2010) to estimate the effect size (i.e., fold change [log_2_], log_2_FC) and sign (i.e., depleted or enriched) of each differentially abundant OTU attributed to individual predictors and their interactions, as well as between different levels within a predictor. Similar to mvabund, edgeR also uses GLMs with negative binomial errors, but it models individual OTU abundances with false discovery rate control (FDR; alpha = 0.05) for multiple testing. Here we referred to edgeR GLMs as FDR-GLMs. FDR-GLMs allow a design matrix accommodating complex experimental structure. Our design matrix followed PC1.prism + PC2.prism + 0 + Group, in which Group contained all combinations of different levels of predictors (Species, Sex and Organ). The model was fitted using glmQLFit() and specific contrasts were made by glmQLFTest(). The *P* values were adjusted by the Benjamini–Hochberg (BH) correction using p.adjust(). We also conducted differential analyses for organ-specific microbial communities to detect differentially abundant OTUs between species within each organ type.

## RESULTS

### Root, leaf and flower microbiomes of *Fragaria*

We analyzed prokaryotic microbiomes associated with roots, leaves and flowers in *F. chiloensis* (F.chilo), *F. virginiana* ssp. *platypetala* (F.virg) and *F. ×ananassa* ssp. *cuneifolia* (F.cunei). These perennial plants and associated microbiota experienced distinct abiotic environments in their native ranges (Figure 1B), and PC1 and PC2 can account for 92% of the abiotic variation. Amplicon sequencing of 16S rRNA with *Arabidopsis*-derived PNA clamps generated substantial chloroplast and mitochondrial sequences (averagely 63% of total reads per sample). After removing the plant contaminants, low-frequency OTUs and low-coverage samples, OTU observations averaged 7896 per sample (median = 2796, s.e. = 1172). Individual microbial communities were normalized (see Methods), and the resultant microbial community matrix consisted of 1577 OTUs from 27 root, 22 leaf and 23 flower samples of the three host species. These OTUs represented the core set of microbiota in our samples.

The root, leaf and flower microbiomes contained predominantly bacterial taxa (with only one Archaea OTU), among which Proteobacteria, Bacteroidetes and Actinobacteria were the most abundant phyla (Figure 1C–E). At the OTU level, root microbiomes possessed more OTUs than leaf and flower microbiomes (Figure 1F; Figure S1). The majority of leaf (94%) and flower (98%) OTUs were also found in roots (Figure 1F). By contrast, root microbiomes harbored more unique OTUs, with only 40% of the OTUs shared with leaf microbiomes, and 53% with flower microbiomes (Figure 1F).

### Organ type as the primary factor structuring plant-associated microbiota

Organ type predicted species α-diversity of microbial communities (Shannon diversity, *F* = 70.33, df = 2, *P* < 0.001; Figure 2C; Table S1), after controlling for abiotic environments (PC1.clim and PC2.clim), host species and sex in a LMM. Shannon diversity (in terms of LS-mean) was highest in root microbiomes, and significantly decreased in leaf (*t* = 9.20, df = 51, *P* < 0.001) and flower microbiomes (*t* = 10.91, df = 51, *P* < 0.001; Figure 2C). Likewise, root microbiomes harbored significantly higher phylogenetic α-diversity (Figure 2D,E; Table S1), using the metrics (Miller *et al*., 2017) that both scale positively with (e.g., PD; Figure 2D) and are insensitive to species richness (e.g., MPD; Figure 2E). Within AG microbiomes, microbial α-diversity was comparable between leaves and flowers (Figure 2C–E).

**FIGURE 2.**
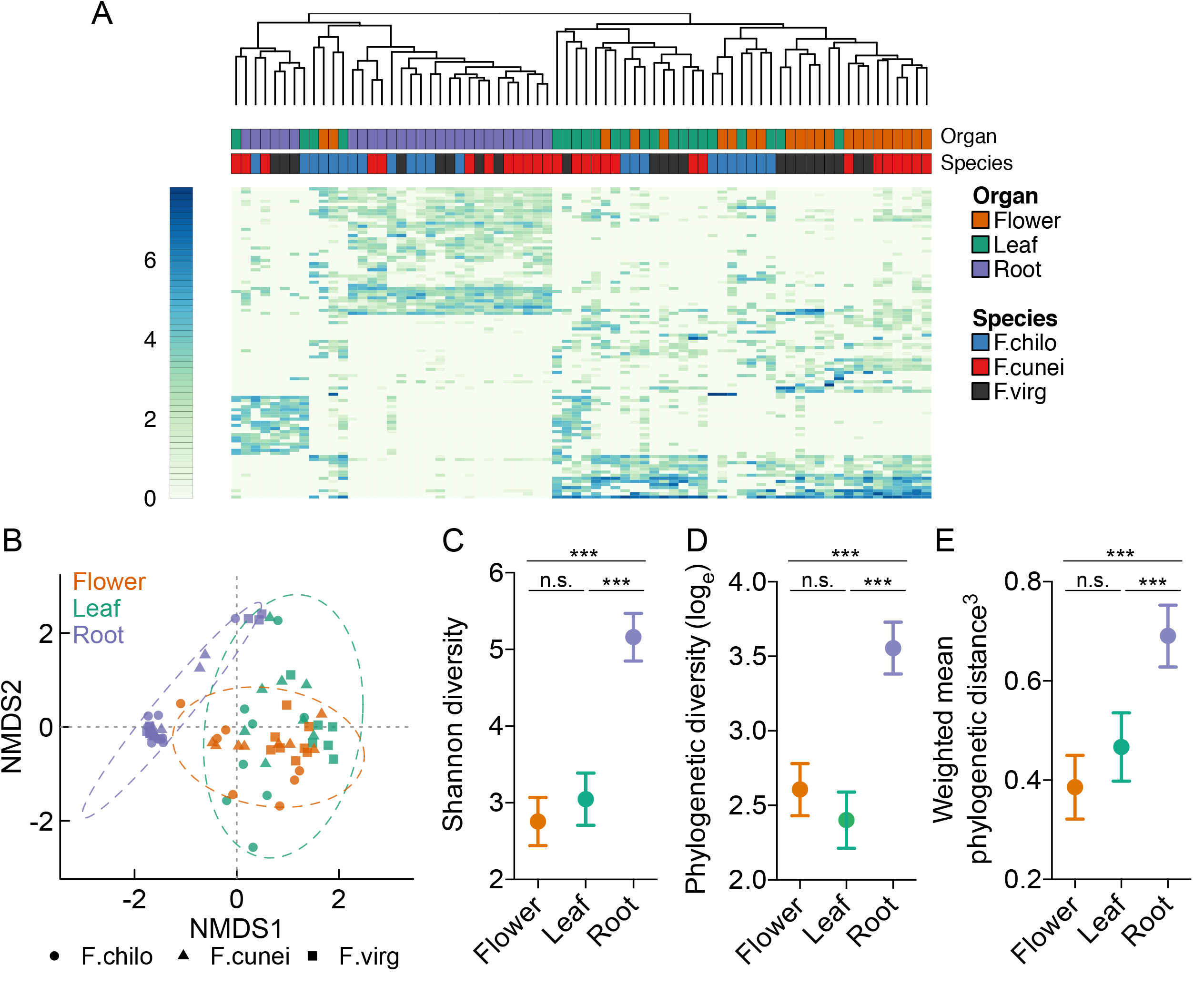
Plants harbor distinct above- and belowground microbiomes. (A) A heat map of the top 100 most abundant OTUs across host species and organ type. The colored scale bar on the left indicates (log_e_) OTU abundances. Hierarchical clustering was performed using the complete linkage method of Euclidean dissimilarity among microbial communities. (B) NMDS of Bray–Curtis dissimilarity revealed that microbial communities were primarily separated according to organ type rather than host species. The ellipses based on 2 s.d. are indicative of the spread of microbial communities within each organ type. (C–E) The least-squares means of Shannon diversity, log-transformed Faith’s phylogenetic diversity (log_e_PD) and power-transformed abundance-weighted mean phylogenetic distance (MPD^3^) are plotted for each organ type, after controlling for all other factors. Error bars represent the 95% confidence intervals. Statistical significance is indicated: ***, *P* ≤ 0.001; n.s., not significant.

Despite substantial overlap in OTUs, AG and BG microbiomes exhibited distinct community structures (Figure 2A,B; Figure S2). Hierarchical clustering analysis (Figure 2A) clearly separated root and leaf/flower microbiomes regardless of host species, and also showed that OTUs of high frequency in roots were not the same ones in leaves and flowers. The NMDS of Bray–Curtis dissimilarity also identified AG and BG organs as the primary source of variation in compositional β-diversity of microbial communities (Figure 2B). Similar to Bray–Curtis NMDS, the separation of root and leaf/flower microbiomes along the dominant ordination axes was revealed by PCoAs of phylogenetic β-diversity metrics (UniFrac and betaMPD; Figure S2).

Consistent with the qualitative inference from NMDS and PCoAs, PERMANOVA showed that organ type accounted for the largest source of variation in microbial β-diversity among the tested predictors (Table S2), when using Bray–Curtis (19.4% of variation, *F* = 7.924, df = 2, *P* = 0.001) and UniFrac distance (21.1%, *F* = 8.838, *P* = 0.001). PERMANOVA of betaMPD agreed with the other two β-diversity metrics that microbial community structures were significantly affected by organ type (*F* = 2.130, df = 2, *P* = 0.001; Table S2), albeit organ type (the main effect) and its interaction with species accounted for a similar amount of variation (6.2% and 7.0%, respectively).

In addition to distance-based PERMANOVA, we used FWER-GLMs to assess community structure changes attributable to each predictor through multivariate modeling of OTU abundances. FWER-GLMs supported the PERMANOVA results that organ type strongly predicted multivariate abundances of microbial communities (deviance = 19,793, df = 2, *P* = 0.001; Table S2). In addition to organ type, host species (deviance = 19,793, *P* = 0.001) and sex (deviance = 12,619, *P* = 0.006) were also identified as having significant impacts on microbial communities, unlike in PERMANOVA (Table S2). The difference between Bray–Curtis PERMANOVA and FWER-GLMs suggested that OTUs responding to host species or sex likely had low or modest variability in abundances between groups of interest, which failed to be detected by less sensitive PERMANOVA (Warton *et al*., 2012), but collectively these OTUs contributed to differential microbial communities. By contrast, OTUs of large among-organ variability were likely involved in distinguishing AG and BG microbiomes, as detected by both PERMANOVA and FWER-GLMs.

FWER-GLMs that detected significant microbial community differentiation caused by organ type also identified the responsible OTUs (Table S3). These differentially abundant OTUs (*N* = 120) attributable to organ type accounted for a relatively small portion (8%) of the overall OTUs (Figure 3; Table S3). We further assessed the effect size and sign of differentially abundant OTUs in leaf or flower microbiomes (relative to root microbiomes), controlling for all other factors using FDR-GLMs. As a result, 414 OTUs were identified as differentially abundant between leaf and root microbiomes, and 404 OTUs between flower and root microbiomes (Figure 3; Figure S3; Table S3), more than three times the OTUs identified by FWER-GLMs. This distinction was primarily caused by stringent FWER control in mvabund GLMs compared to FDR control in edgeR GLMs.

**FIGURE 3.**
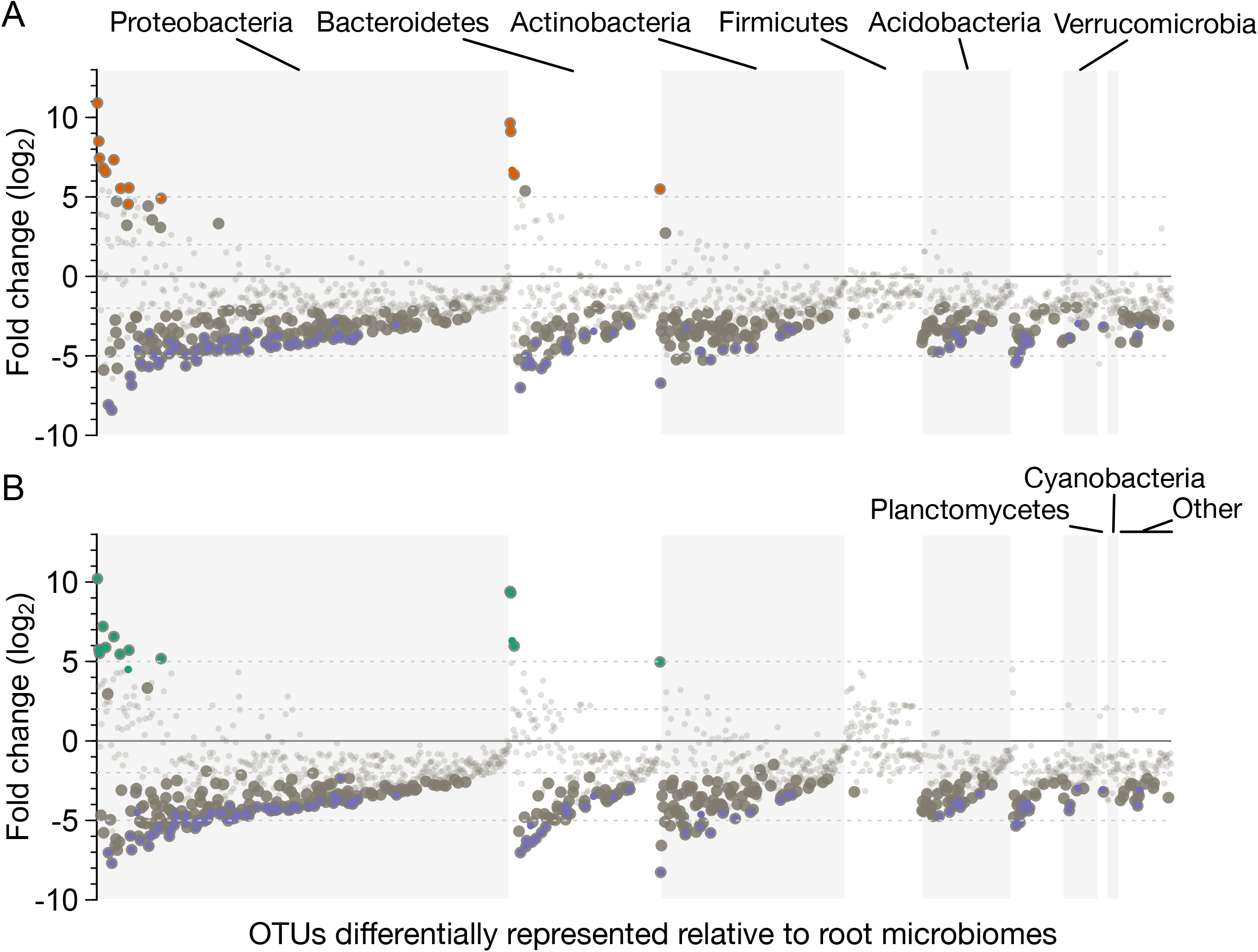
Differentially abundant OTUs in flower (A) and leaf (B) relative to root microbiomes. (A) Significantly enriched and depleted OTUs in flower relative to root microbiomes are shown as large, dark gray dots, identified by FDR-GLMs. OTUs with non-significant abundance changes are shown as small, light gray dots. The orange and purple dots of the immediate point size denote significant OTUs of differential abundances in response to organ type, identified by FWER-GLMs. (b) FDR-GLMs were also conducted to identity enriched and depleted OTUs in leaf relative to root microbiomes. The green and purple dots denote the OTUs responding significantly to organ type, identified by FWER-GLMs. In both panels, bacterial phyla and individual OTUs within each phylum are ordered along the *x*-axis by their relative abundances in the whole data set. Dashed horizontal lines depict (log_2_) fold change of 2 and 5 (as well as −2 and −5).

Depletion effects dominated AG microbiomes relative to BG microbiomes (Figure 3). In flower microbiomes, FDR-GLMs identified 22 OTUs as significantly enriched and 382 OTUs as significantly depleted (Figure 3A, large gray dots). The enriched OTUs concentrated in the three dominant phyla. Thirteen out of the 22 enriched OTUs in flower microbiomes, with large effect size (typically of≥5 log_2_FC), overlapped those identified by FWER-GLMs as responsive to organ type (Figure 3A, orange dots). These large-effect, enriched OTUs were primarily from *Sphingomonas, Hymenobacter, Janthinobacterium, Pseudomonas, Methylobacterium* and *Salinibacterium* (Table S3). The 382 OTUs that were significantly depleted from flower microbiomes ranged across diverse phyla (Figure 3A; Table S3), among which 95 OTUs were also identified by FWER-GLMs as significantly influenced by organ type (Figure 3A, purple dots). These 95 OTUs had (log_2_) fold changes averaged −4.50 (range −2.92 to −8.42; Table S3), among which OTUs from *Streptomyces, Bradyrhizobium* and *Steroidobacter* showed the largest effect sizes (in absolute values).

Flower and leaf microbiomes were similar in enriched and depleted OTUs in several ways (Figure 3; Figure S3; Table S3). First, more than 90% of the 414 differentially abundant OTUs identified by FDR-GLMs in leaf microbiomes overlapped those detected in flower microbiomes (Figure S3). Second, these overlapping OTUs between leaf and flower microbiomes were also correlated in their fold changes (Pearson's *r* = 0.947, *t* = 58.05, df = 384, *P* < 0.001). Third, leaf and flower microbiomes had the same set of FDR-GLM detected, enriched (except one OTU) and depleted OTUs, overlapping those identified by FWER-GLMs as responsive to organ type (Figure 3B, colored dots; Table S3).

### Increased host species influence from root to leaf to flower microbiomes

As plants harbored distinct AG and BG microbiomes, host species effect (see definition in Introduction) was assessed for root, leaf and flower microbiomes separately, while controlling for abiotic environments and host sex. Host species did not predict microbial α-diversity for any organ-associated microbiomes, nor did abiotic environments (Table 1). Different from α-diversity measures, the extent to which host species overlapped in OTUs changed from BG to AG microbiomes (Figure 4A–C). In root microbiomes, the three *Fragaria* species overlapped substantially in OTUs, accounting for 71–82% of individual host species OTUs (Figure 4A); but this OTU sharing dropped to 20–58% in leaf and 27–45% in flower microbiomes (Figure 4B,C), suggesting that microbial communities could be more similar among host species in roots, compared to leaves and flowers.

**FIGURE 4.**
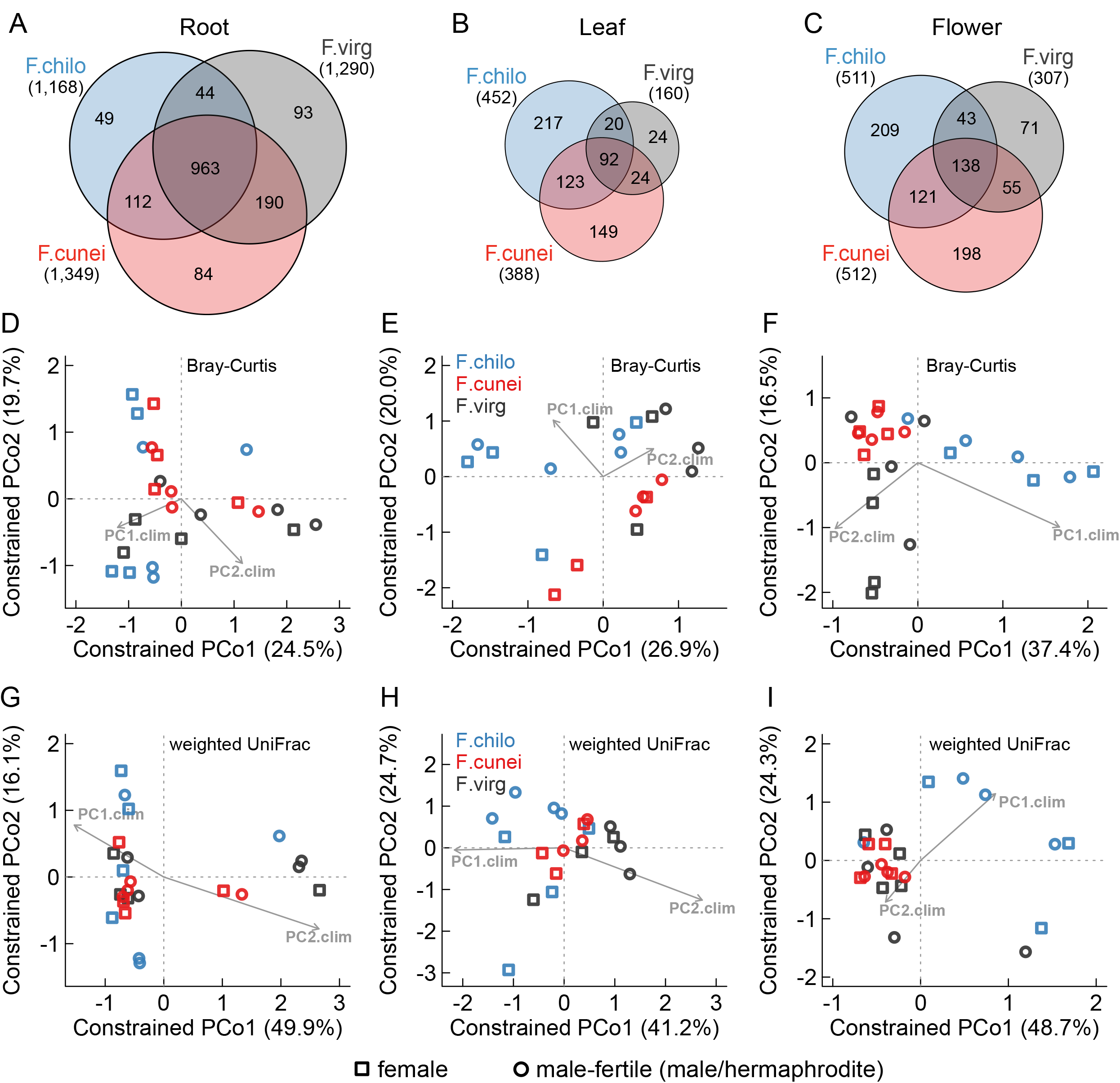
Host plant species influences aboveground but not belowground microbiomes. (A–C) OTU overlap among the three host species for root, leaf and flower microbiomes, respectively. (D–F) Constrained PCoAs of Bray–Curtis dissimilarity indicated enhanced microbial community separation by host plant species from root (D) to leaf (E) and flower (F) microbiomes, controlling for abiotic environments (PC1.clim and PC2.clim) and sex (female and male/hermaphrodite). (G–I) Constrained PCoAs of abundance-weighted UniFrac distance for root, leaf and flower microbiomes, respectively.

**TABLE 1.**
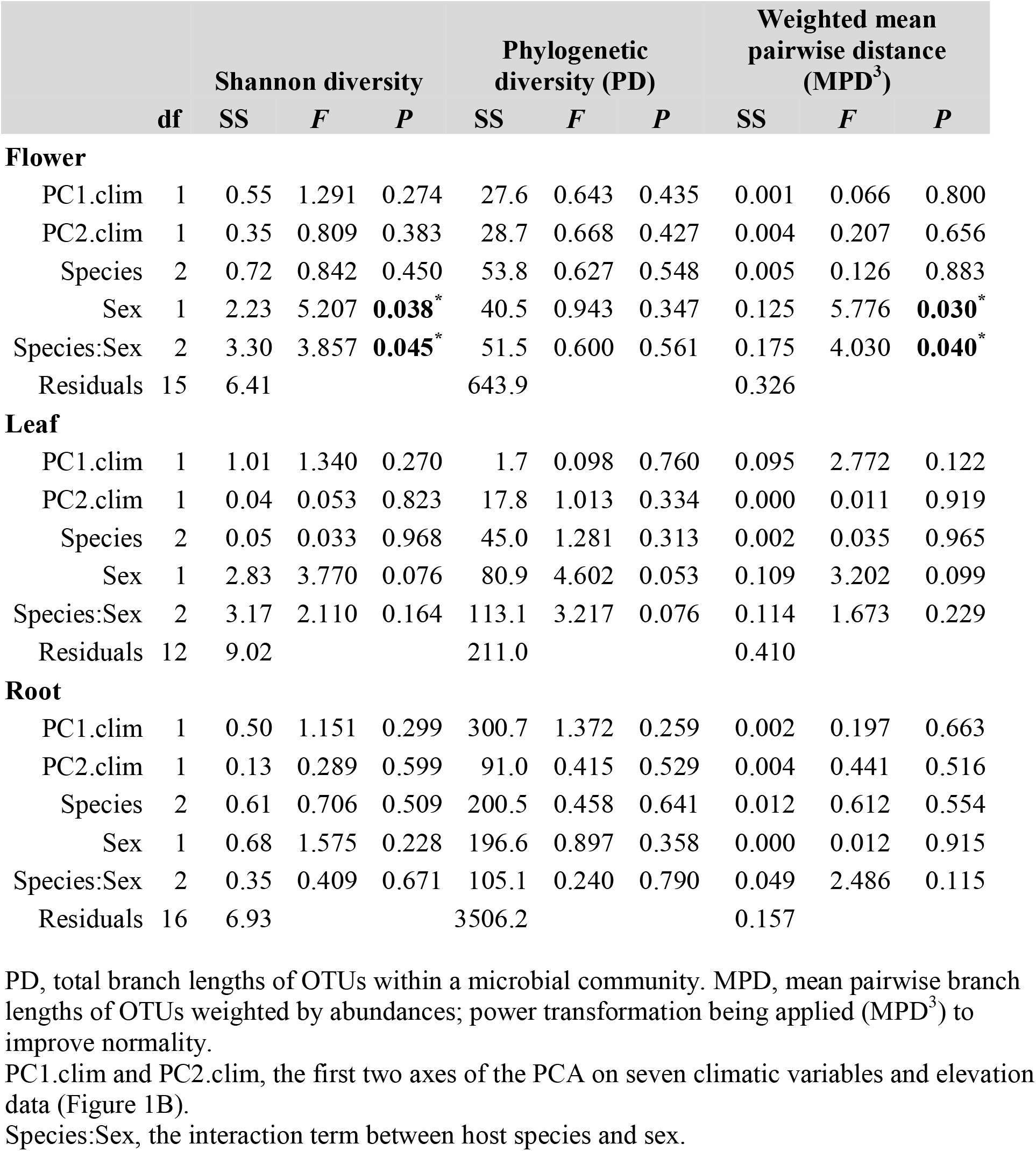
General linear models testing the responses of species and phylogenetic α-diversity of microbial communities to abiotic environments, host species and host sex. Statistical significance was evaluated by Type III sums of squares (SS).

In support of this inference, constrained PCoAs of compositional and phylogenetic β-diversity among organ-specific microbial communities (Figure 4D–I; Figure S4) detected diminishing community separation by host species from AG to BG microbiomes, while controlling for all other factors. For flower microbiomes (Figure 4F,I; Figure S4), F.chilo-associated microbial communities were segregated with the other two species along the first axis of constrained PCoAs for all three β-diversity metrics, whereas those of F.virg and the hybrid F.cunei were separated by the second axis for compositional β-diversity (Bray–Curtis) metric. In leaf microbiomes, community separation was only seen when using Bray–Curtis metric between the hybrid F.cunei and two parental species along the second axis (Figure 4E); when using phylogenetic β-diversity metrics, host species overlapped in microbial communities (Figure 4H; Figure S4). This microbiome overlapping among host species was most pronounced in roots.

Consistent with the constrained PCoAs results, PERMANOVA (Table S4) showed that host species explained the largest source of variation in flower microbiomes among all tested predictors (Bray–Curtis, 12.9% of variation, df = 2, *F* = 1.507, *P* = 0.049; UniFrac, 20.6%, *F* = 2.920, *P* = 0.004; betaMPD, 13.1%, *F* = 1.575, *P* = 0.006). Further evidence came from FWER-GLMs that host species strongly predicted multivariate abundances of flower microbiomes (deviance = 2120, df = 2, *P* = 0.030; Table S4), while controlling for all other factors. In leaf microbiomes, UniFrac and betaMPD PERMANOVA corroborated the inference from constrained PCoAs that host species did not affect the phylogenetic community structures of leaf microbiota (F = 0.827, df = 2, *P* = 0.6 and *F* = 1.001, *P* = 0.4, respectively; Table S4). Yet, the subtler microbial community separation by host species in leaves relative to flowers (Figure 4E,F) was captured by FWER-GLMs (deviance = 1771, df = 2, *P* = 0.031; Table S4), albeit not by less sensitive PERMANOVA (Bray–Curtis, *F* = 0.966, df = 2, *P* = 0.5). By contrast, in root microbiomes both FWER-GLMs and Bray–Curtis PERMANOVA (Table S4) supported that host species did not predict multivariate abundances of root microbial communities (deviance = 4657, df = 2, *P* = 0.056 and *F* = 1.131, *P* = 0.3, respectively), as well as the phylogenetic community structures (UniFrac PERMANOVA, *F* = 0.887, *P* = 0.5; betaMPD, *F* = 0.959, *P* = 0.8). Across root, leaf and flower microbiomes, abiotic environments (PC1.clim and PC2.clim together) explained a similar amount of variation in microbial community structures as did host species (PERMANOVA, Table S4).

FWER-GLMs and FDR-GLMs were used to identify the OTUs underlying microbial community differentiation caused by host species, for flower and leaf microbiomes separately. Surprisingly, no OTUs were reported as differentially abundant among host species in flowers or leaves (Tables S5, S6). However, the effect size estimation by FDR-GLMs indicated that many OTUs exhibited relatively large fold changes (log_2_FC in absolute value ≥ 5) among host species, accounting for 37% of flower and 53% of leaf OTUs (Tables S5, S6); this was likely attributable to a great deal of host species-specific OTUs and limited OTU overlapping among all three host species (Figure 4B,C). The presence of non-significant, large-effect OTUs also suggested considerable variation in OTU abundances within host species given our small sample sizes, which perhaps limited the power to detecting OTU-level but not yet community-level differences among host species.

### Sexual dimorphism in microbiomes contingent upon host species and organ type

Sexual dimorphism was assessed for root, leaf and flower microbiomes separately. In LMs (Table 1), the main effect of sex informed whether microbial communities responded to sex type in general (i.e., averaged across species), whereas its interaction with species indicated whether such responses were host species specific. We found that sex predicted species and phylogenetic α-diversity of microbial communities in flowers, but not in leaves and roots (Table 1). In flower microbiomes, sex comprised the second largest source of variation in Shannon diversity (16.4% of variation, *F* = 5.207, df = 1, *P* = 0.038; Table 1) and MPD (19.7%, *F* = 5.776, *P* = 0.030), although not in PD (4.8%, *F* = 0.943, *P* = 0.3). Compared to the main sex effect, species-specific sex effect on flower microbial α-diversity was even stronger, explaining the largest source of variation (Shannon diversity, 24.3%, *F* = 3.857, df = 2, *P* = 0.045; MPD, 27.5%, *F* = 4.030, *P* = 0.040). Specifically, males harbored higher microbial α-diversity than females in F.chilo (Shannon diversity, *F* = 5.207, df = 1, *P* = 0.038; MPD, *F* = 5.776, *P* = 0.030; Figure S5) that also has the most pronounced sexual dimorphism (see Discussion); but in the other two species, inter-sexual differences were not significant (Figure S5).

In contrast, microbial community β-diversity was not influenced by the main effect of sex (averaged across species), while controlling for all other factors (Table S4). This pattern was consistent across organ types, β-diversity metrics and statistical models (PERMANOVA and FWER-GLMs). However, species-specific sex effects on community structures were observed in flower and leaf microbiomes by FWER-GLMs (Table S4), although such interaction effects failed to be captured by less sensitive PERMANOVA.

Because small sample sizes limited the detectability of differentially abundant OTUs, we focused on phylum-level variation in relative abundances between intraspecific male/hermaphrodite and female hosts for flower and leaf microbiomes. In flowers (Figure S6), males/hermaphrodites harbored proportionately more Bacteroidetes (in all three host species, *P* < 0.001 in proportion tests) and less Proteobacteria (all *P* < 0.001) than females, after controlling for FDR (alpha = 0.05). For flower Actinobacteria, the relative abundances were also higher in hermaphrodites but only in two species (F.chilo and F.cunei, *P* < 0.001). In leaves (Figure S7), sex differences in the three dominant phyla were variable among host species.

## DISCUSSION

By characterizing the microbiomes of the three wild strawberry species *in situ*, we quantified the respective effects of organ type, host species, and host sex on shaping these plant-associated microbiota in their wild populations. Our modeling of microbial α- and β-diversity revealed organ type as the primary factor structuring microbial communities, exceeding the effect of host species, even in the wild. Moreover, by comparing host influence on organ-specific microbial communities, we found enhanced host species effect from root to leaf to flower microbiomes. Lastly, our study presented the first evidence of sexual dimorphism in leaf and flower bacterial communities, and we found that such sexual dimorphism was contingent upon host species.

### Organ type structures plant-associated microbiota in the wild

Our data support the hypothesis that organ type has a preeminent role in structuring plant-associated microbiomes across host species (Vorholt, 2012; Müller *et al*., 2016). In the three wild strawberries in their native environments, organ type not only predicts species and phylogenetic α-diversity of plant-associated microbiomes, but also explains the largest source of variation in compositional and phylogenetic β-diversity, while controlling for the effects of their broad-scale abiotic environments, host species and sex. In other words, the root microbiome of one host species is expected to be more similar to that of a different host species than to its own leaf and/or flower microbiome, potentially owing to niche-specific selection for adapted microbiota in different plant organs (Vorholt, 2012; Bai *et al*., 2015; Müller *et al*., 2016).

In line with earlier studies on root microbiomes (e.g., Schlaeppi *et al*., 2014), Proteobacteria (45%, relative abundance), Actinobacteria (22%) and Bacteroidetes (14%) are the dominant bacterial phyla in *Fragaria*, with Firmicutes to a lesser extent (6%). However, these previous studies often identified substantial influence of soil type on root microbiomes in both greenhouse and manipulated field settings, stronger than the effects of host genotypes (Bulgarelli *et al*., 2012; Edwards *et al*., 2015) and plant species (Schlaeppi *et al*., 2014). By contrast, our study observed convergence in root microbiomes, despite distinct soil habitats where the three *Fragaria* species grow (Salamone *et al*., 2013).

The discrepancy with previous studies on the relation of root microbiomes with soil habitats likely has two reasons. First, root microbiomes quantified here comprise microbiota in association with absorptive fine roots (or first-order roots), as compared to those associated, for example, with the primary root (Bulgarelli *et al*., 2012; Schlaeppi *et al*., 2014) or the whole root system (Lundberg *et al*., 2012; Edwards *et al*., 2015) in *Arabidopsis* and rice. It is possible that metabolically active root parts impose stronger filtering for colonizing microbes and thus cause stronger deviation from source soil microbiota, relative to metabolically inactive parts. Nevertheless, this hypothesis requires further investigation relating root traits to root microbiomes. In fact, several root samples in this study comprising old segments of fine roots, which were perhaps no longer metabolically active, formed a separate cluster different from the other root samples (Figure 2A). Second, root microbiomes here did not consider low-frequency OTUs, owing to sequencing depth constraint and the use of normalized microbial community matrix and abundance-weighted diversity indexes. Although we cannot rule out the possibility that the low-frequency OTUs may indeed differ with soil habitats, their influence on microbial community structure and function remains an open question.

*Fragaria* leaf and flower microbiomes shared most of their OTUs with root microbiomes. This is in line with the idea that the sources of microbial assemblies in phyllosphere (Vorholt, 2012) involve colonizing microbiota from soil by rain splash, wind and the visits of ground-dwelling herbivores and pollinators, as well as endophytes migrating from root to AG organs (Chi *et al*., 2005). Consistent with grapevine AG and BG microbiomes in managed field settings (Zarraonaindia *et al*., 2015), we also found that leaf and flower microbiomes shared proportionally more OTUs with root microbiomes than with each other, indicating soil microbiota as a common species pool for microbial assemblies associated with different plant organs. Despite substantial OTU overlap, AG and BG microbiomes differed significantly in community structures, as has been observed in leaf and root microbiomes of *Arabidopsis* in both controlled and wild settings (Bodenhausen *et al*., 2013; Bai *et al*., 2015). Such AG–BG microbiota differentiation in *Fragaria* was attributable to many depleted bacterial taxa in phyllosphere, especially *Bradyrhizobium* and *Steroidobacter*. These two bacterial genera were also found enriched in grapevine root microbiome (relative to AG microbiomes), where they probably mediate essential processes including nitrogen fixation in roots in vineyards (Zarraonaindia *et al*., 2015). The enriched taxa in *Fragaria* leaves and flowers such as *Methylobacterium, Sphingomonas* and *Pseudomonas* have been detected in other plant species, and their abilities to withstand more hostile habitats of phyllosphere have been implicated (reviewed in Vorholt, 2012; Aleklett *et al*., 2014).

### Host species influence increases from BG to AG microbiomes

Although in wild populations plant species–microbiota association can be influenced by abiotic environments, our study showed that host species differences in microbiomes were not just the by-product of their broad-scale variation in abiotic environments, but that host species explained a substantial amount of variation of organ-specific microbial communities, after accounting for that explained by abiotic environments.

The effect of *Fragaria* host species was strongest in flower microbiomes followed by leaf, and weakest in root microbiomes. Such enhanced host influence from root to leaf microbiomes has been detected in other plants at both genotype (Wagner *et al*., 2016) and species levels (David *et al*., 2016). In common garden experiments of *Boechera stricta*, phylogenetic community structure of leaf bacterial microbiota was clustered in accordance with host genotype but not for root microbiota (Wagner *et al*., 2016). Similar to our finding of *Fragaria* species influence on bacterial microbiomes, host species was found as significantly affecting fungal endophytic communities in leaves, but not affecting those in roots, among three common grass species in their wild populations (David *et al*., 2016). If these patterns hold true across plant taxa, we may expect relaxed phylogenetic conservatism of plant microbiome traits (e.g., community structure) from BG to AG, owing to the paralleling phylogenetic signal of host plant functional traits.

Studies examining root and leaf traits (Valverde-Barrantes *et al*., 2015; Weemstra *et al*., 2016) have revealed stronger phylogenetic conservatism in roots, with root trait variation largely among deeply diverged plant lineages and little variation among species of low taxonomic ranks. The three *Fragaria* congeners in this study have a relatively short divergence history (originated ~1 Mya) and identical creeping herbaceous life form (Liston *et al*., 2014), and thus they likely have similar fine root traits. This perhaps explains the resemblance of root microbiomes among host species in our study. Compared to root traits, interspecific differentiation in leaf traits (e.g., coriaceousness, leaf thickness) have been seen among the three *Fragaria* species in the wild (Figure 1A) and greenhouse (Salamone *et al*., 2013); this may underlie the more heterogeneous leaf microbiomes relative to root microbiomes among host species. Relative to root and leaf traits, floral traits (e.g., color, size, scent and reward) are exceedingly diverse in plants (Stebbins, 1970; Fenster *et al*., 2004), and thus flower microbiomes are predicted to be distinct among host species (Aleklett *et al*., 2014). Interestingly, although *Fragaria* flowers are morphologically similar in shape and color (Figure 1A; see also Liston *et al*., 2014), microbiome divergence among host species was yet strongest in flowers, suggesting that other floral traits such as scent, and pollen and nectar rewards might be critical in shaping species-specific flower microbiome. For plant genera with highly diversified floral traits, host species influence on flower microbiomes may be even stronger than what we observed in *Fragaria*.

### Sexual dimorphism is present in flower and leaf microbiomes

Intriguingly, we found that the degree of sexual dimorphism in microbiomes coincided with the degree of sexual dimorphism in the three host species. In wild strawberries, *F. chiloensis* has the most complete separation of sexes (dioecy) than other *Fragaria* species, and has perhaps the greatest sexual differentiation in floral and other traits (Ashman *et al*., 2012). In *F. chiloensis*, male flowers, for example, are typically larger in petal size than female flowers (Ashman *et al*., 2012), which likely in part explains higher species and phylogenetic α-diversity of flower microbiomes in males. Although comparable studies on sexual dimorphism in plant microbiomes are lacking, it is noteworthy that culturable nectar-dwelling yeasts appeared to be higher in richness in male than female flowers of *Silene latifolia* in the wild (Golonka and Vilgalys, 2013).

Relative to α-diversity, microbial community structure seems more sensitive to host sex, as many bacterial phyla were found differentially abundant between sexes in flower and leaf microbiomes across species. Sexually dimorphic leaf traits, which may underlie the observed differences in leaf microbiomes between sexes, have been detected in *F. chiloensis* and *F. virginiana* ssp. *platypetala* in common gardens (T-L Ashman and N Wei, unpubl. res.). For both species, males/hermaphrodites possess higher leaf nitrogen content and specific leaf area than females; but the degree of sexual dimorphism in leaf traits is still higher in *F. chiloensis* than *F. virginiana* ssp. *platypetala*. Although similar data are not available for the hybrid *F. ×ananassa* ssp. *cuneifolia*, we suspect that it can be similar to *F. virginiana* ssp. *platypetala* considering their morphological resemblance (Salamone *et al*., 2013). Nevertheless, exactly how these sexually dimorphic traits affect microbiomes is not clear, but deserves additional research.

To conclude, our study provides the first characterization of microbiomes associated with the close wild relatives of the cultivated strawberry. We show, for the first time, enhanced host species influence and sexual dimorphism from root to flower microbiomes in wild populations. While these findings await similar investigations to generalize how plants control microbiota assemblies in the wild, it is important to recognize that such patterns of host species–microbiota association *in situ* affect plant interactions with AG and BG environments and plant fitness. Moreover, our results of sex-differential microbiota expand the understanding of sexual dimorphism in plants, and also highlight the needs for future research on the underlying mechanisms and on relating these differences to sex-specific fitness. Overall, findings from the wild, like ours here, strengthen those from experimental settings, and together they have broad implications for understanding this extended phenotype of plants.

## AUTHOR CONTRIBUTIONS

N.W. and T-L.A. designed research and conducted sampling; N.W. performed research and analyzed data; N.W. and T-L.A. wrote the paper.

## ACKNOWLEDGEMENTS

We thank Chris Marshall for generous help with QIIME pipeline, Aaron Liston for providing population locations and Anna Freundlich for assistance with sampling, and the Ashman lab members for discussion.

